# Evolutionary trajectories of immune escape across cancers

**DOI:** 10.1101/2025.01.17.632799

**Authors:** Wenjie Chen, Toby Baker, Zhihui Zhang, Huw A. Ogilvie, Peter Van Loo, Shengqing (Stan) Gu

**Affiliations:** Department of Hematopoietic Biology & Malignancy, The University of Texas MD Anderson Cancer Center, Houston, Texas, USA; Department of Genetics, The University of Texas MD Anderson Cancer Center, Houston, Texas, USA; The Francis Crick Institute, London, United Kingdom; Department of Medicine, University of California Los Angeles, Los Angeles, California, USA; Department of Genomic Medicine, The University of Texas MD Anderson Cancer Center, Houston, Texas, USA

**Keywords:** Immune escape, cancer evolution, mutation timing

## Abstract

Immune escape is a critical hallmark of cancer progression and underlies resistance to multiple immunotherapies. However, it remains unclear when the genetic events associated with immune escape occur during cancer development. Here, we integrate functional genomics studies of immunomodulatory genes with a tumor evolution reconstruction approach to infer the evolution of immune escape across 38 cancer types from the Pan-Cancer Analysis of Whole Genomes dataset. Different cancers favor mutations in different immunomodulatory pathways. For example, the antigen presentation machinery is highly mutated in colorectal adenocarcinoma, lung squamous cell carcinoma, and chromophobe renal cell carcinoma, and the protein methylation pathway is highly mutated in bladder transitional cell carcinoma and lung adenocarcinoma. We also observe different timing patterns in multiple immunomodulatory pathways. For instance, mutations impacting genes involved in cellular amino acid metabolism were more likely to happen late in pancreatic adenocarcinoma. Mutations in the glucocorticoid receptor regulatory network pathway tended to occur early, while mutations in the TNF pathways were more likely to occur late in B-cell non-Hodgkin lymphoma. Mutations in the NOD1/2 signaling pathway and DNA binding transcription factor activity tended to happen late in breast adenocarcinoma and ovarian adenocarcinoma. Together, these results delineate the evolutionary trajectories of immune escape in different cancer types and highlight opportunities for improved immunotherapy of cancer.

**Significance:** Despite its critical role in cancer progression, the evolution of immune escape is poorly understood. We integrate functional genomics and tumor evolution reconstruction and infer immune escape trajectories across cancer types. Our results have important implications for developing biomarkers for immunoprevention and treatment strategies for immune escape of cancer.

## INTRODUCTION

Immune selection is a major driver of cancer evolution (1,2), and immune escape is a critical hallmark of cancer progression (3,4). Various immune escape mechanisms have been discovered (5,6), among which impairment of antigen presentation is a frequent event (7–9). However, it remains unclear when the genetic events that underlie immune escape occur during cancer development. Gaining such knowledge can help us understand how immune resistance emerges, what genetic events precede its emergence, and how cancers further evolve after acquiring immune resistance. This in turn can help devise personalized therapeutic strategies that block the adaptation of cancers to evolving selection pressures.

As our understanding of anti-cancer immunity deepens, the evolution of immune escape has elicited intense interest in the field of cancer immunology. Multiple studies performed multi-region sampling (such as primary and metastatic sites) followed by whole exome/genome sequencing to infer the evolution of immune escape in each patient (10–14). These studies have led to recurrent observations of impairment of (neo)antigen presentation in early-stage tumors (11,15). However, this approach relies heavily on the availability of multi-region samples and therefore sample size has been limited. Besides, this approach cannot compare the timing of clonal mutations that are present in all cancer cells in the tumor. Several studies also delineated the genetic landscape that underlies immune escape in primary and metastatic cancers (16,17). For example, a recent study characterized mutations in six molecules or pathways that can cause immune escape: HLA, antigen presentation, IFNγ pathway, PD-L1 amplification, CD58 alteration, and epigenetic regulation by SETDB1 (16). Nevertheless, the selection of immune escape pathways in existing studies is often limited and arbitrary, with a bias toward well-documented pathways such as antigen presentation.

In this study, we utilize a tumor evolution reconstruction approach and leverage our recently developed GRITIC tool (18) to perform fine-scaled timing of events in tumor evolution. We also utilize published functional genomics screens to systematically identify the immunomodulatory genes and pathways whose mutation might underlie immune escape. We first focused on the antigen presentation machinery (APM) and inferred the timing of mutations in genes involved in this pathway. Next, we expanded to the core pathways that regulate other aspects of cancer-immune interaction and delineate the evolutionary trajectories of mutations in these pathways.

## RESULTS

### Prevalence of antigen presentation machinery mutations across cancer types

The antigen presentation machinery (APM) mediates the recognition of cancer cells by T cells. Its impairment is prevalent in cancer and is a major mechanism of resistance to immunotherapies. Antigen presentation mediated by Class I Major Histocompatibility Complex (MHC-I) is prevalent in all cell types, whereas antigen presentation mediated by Class II MHC (MHC-II) is mostly present in antigen presenting cells (APCs) (19). Therefore, we focused on mutations in the MHC-I mediated APM. We first examined the prevalence of APM mutations in the Pan-Cancer Analysis of Whole Genomes (PCAWG) dataset with 2,658 whole genome sequences (WGS) of cancers from 38 cancer types (**Fig. S1A**). 7.7% of samples showed point mutations in the APM, dominated by *NLRC5* and *B2M* mutations (**Fig. 1A**). In addition, 18.9% of samples showed HLA loss of heterozygosity (LOH) (**Fig. 1A**). Multiple cancer types showed a high prevalence of APM mutations, such as chromophobe renal cell carcinoma, lung squamous cell carcinoma, B-cell non-Hodgkin lymphoma, and colorectal adenocarcinoma (**Fig. 1B** and **S1B**). These findings align with previous studies reporting APM mutations in various cancers (10,16,20).

**Figure 1.**
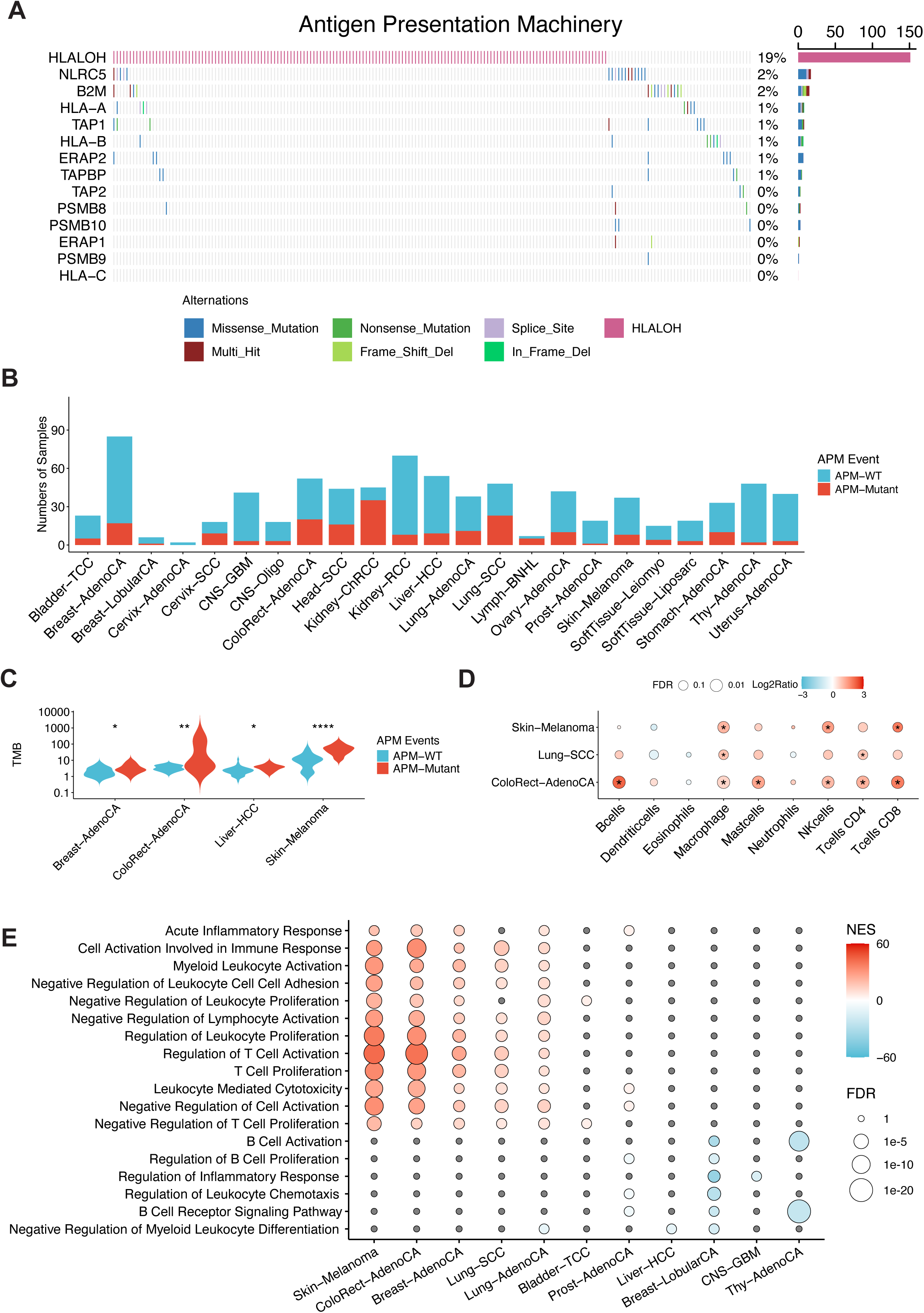
Genetics of antigen presentation machinery across cancer types. (A) Prevalence of antigen presentation machinery mutations in PCAWG-TCGA samples. (B) Prevalence of antigen presentation machinery mutations in each PCAWG-TCGA cancer type. (C) Tumor mutation burden with antigen presentation machinery mutation in each cancer type. (D) Different tumor microenvironments inferred by CIBERSORT. The dot size shows the mean ratio of cell score between samples with APM and the ones without APM mutations. The color shows the FDR for adjusting every comparison using the Wilcoxon test. * show the significance of the comparison. (E) Differential enrichments by antigen presentation machinery mutations in multiple cancer types. The red box presents the up-regulation of the gene set in the samples with APM mutations and the blue box shows the down-regulated ones. The gray box indicates insignificant enrichments.

Tumor mutation burden (TMB) is often associated with immune selection pressure. Indeed, we found in multiple cancer types that tumor samples with APM mutation showed higher TMB levels (**Fig. 1C**). Furthermore, compared with tumors without APM mutation, those with APM mutation showed higher levels of immune infiltration in multiple cancer types, including melanoma, lung squamous cell carcinoma, and colorectal adenocarcinoma (**Fig. 1D**). Comparison of bulk tumor gene expression showed that in breast adenocarcinoma, colorectal adenocarcinoma, lung cancer, and melanoma, samples with APM mutation showed higher expression of genes involved in inflammation and T cell activation (**Fig. 1E**). Conversely, in breast lobular carcinoma, thyroid adenocarcinoma, and prostate adenocarcinoma, samples with APM mutation showed lower expression of genes involved in B-cell-mediated immune response (**Fig. 1E**).

Next, we investigated whether APM point mutations are under selection across cancer types using the non-synonymous:synonymous substitution ratio (dN/dS) (21). APM mutations presented a dN/dS ratio of >1 in truncating mutations in a pan-cancer analysis. Positive selection of truncated mutations in APM was also found in colorectal adenocarcinoma and B-cell non-Hodgkin lymphoma (**Fig. S1C**).

### Timing of mutations in the antigen presentation machinery across cancer types

We then inferred the timing of mutations in the APM across cancer types in all PCAWG samples leveraging the principles of GRITIC (18) to time individual mutations, and characterized how cancers evolve before or after acquiring mutations in the APM. Briefly, our approach uses read counts to infer the relative timing of a given SNV and any overlapping copy number gains. The gain timing distributions from GRITIC are then used to produce posterior timing distributions for each SNV. The SNV timing is on a scale called mutation time which ranges from 0 to 1, with 0 representing conception and 1 representing the end of the tumor’s clonal evolutionary period (18). The relative timing of mutations in each gene was calculated by subtracting the mean timing of all SNVs in driver genes from the gene mutation timing in each sample (**Fig. 2A**).

**Figure 2.**
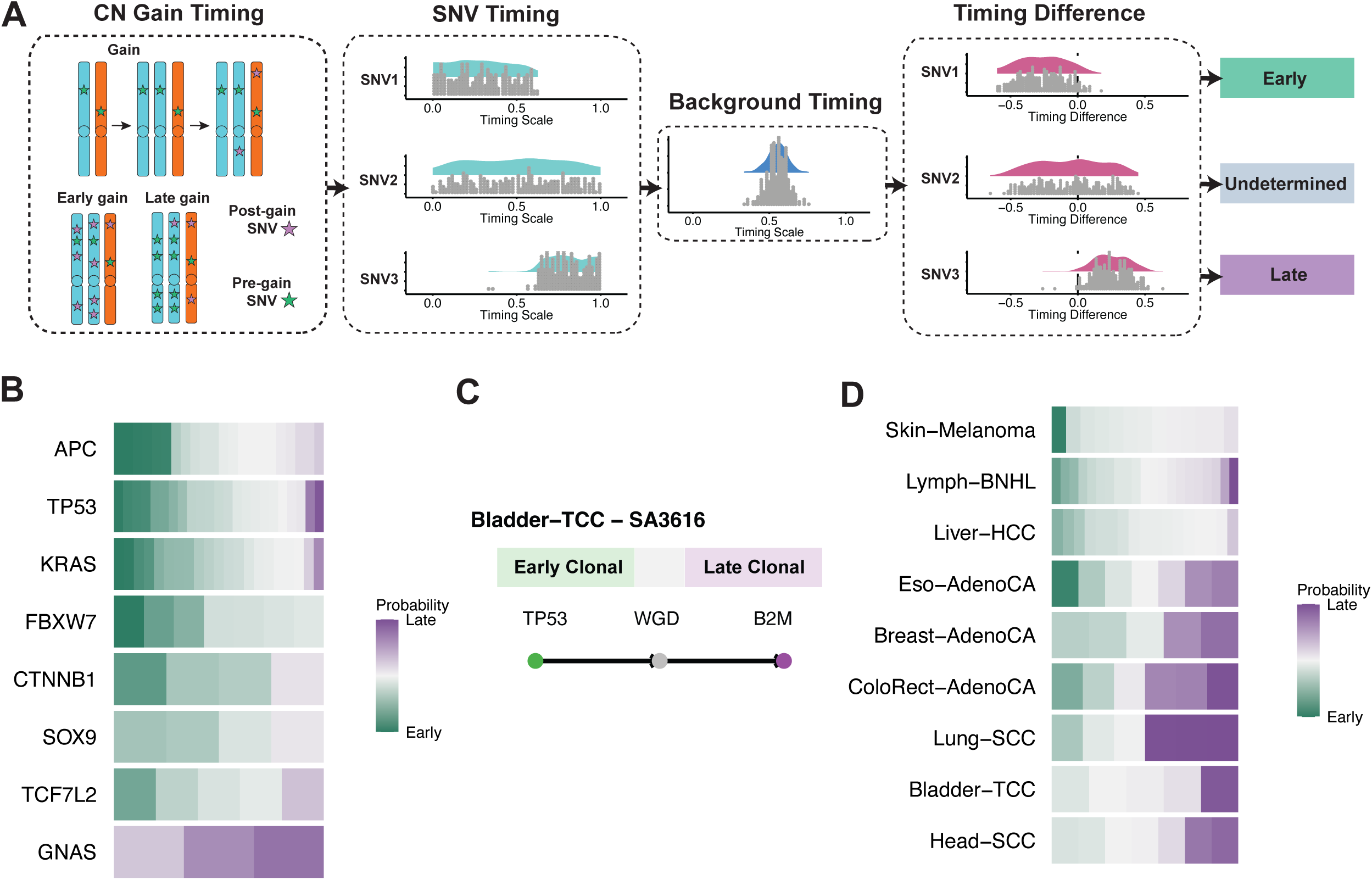
Timing of mutations in the antigen presentation machinery across cancer types. (A) Definition of early/late mutations in cancer samples. CN: Copy number; SNV: single nucleotide variant. (B) Timing of driver genes in stomach adenocarcinoma. Green represents early timing, transitioning gradually to purple for late timing. The color intensity reflects the proportion of early or late events out of 250 samplings, indicating the likelihood of an event occurring early or late. (C) An example of the timing of antigen presentation machinery mutations in bladder transitional cell carcinoma. (D) Comparison of the timing of antigen presentation machinery point mutations and the background timing.

To evaluate the performance of our approach for inferring SNV timing, we first used it to reconstruct event timing in colorectal cancer, where mutations in *APC*, *KRAS*, and TP53 are known to be the early steps in tumor evolution (22–25). Reassuringly, our approach infers that *APC*, *KRAS*, and *TP53* mutations are among the earliest events in colorectal cancer evolution (**Fig. 2B**), consistent with the published studies (22–25).

We then evaluated the timing of APM mutations across cancer types. Of the 137 SNVs in APM genes, 86% were clonal, and 14% were subclonal (**Fig. S2A**). We were able to time most SNVs, particularly clonal SNVs in samples with complex copy number alterations (CNAs), building upon the strengths of GRITIC in timing complex copy number gains (18). For example, in a bladder transitional cell carcinoma patient with clonal mutations in *B2M* and *TP53* and whole genome doubling (WGD), GRITIC inferred that the *B2M* mutation happened after the *TP53* mutation and the WGD (**Fig. 2C**).

We defined the mean timing of all mutations in driver genes as the background timing for each sample. For mutations in the APM, we compared their timing with the background timing and categorized them as “early”, “late”, or “undetermined” in each patient (**Methods**, **Fig. 2A**).

Compared to the background timing in each sample, APM point mutations often happened late in colorectal cancer, squamous cell lung cancer, and head and neck squamous cell carcinoma (**Fig. 2D**).

For samples with HLA LOH, it is difficult to directly infer the timing of LOH, due to the loss of mutation information on the lost allele. As an alternative approach, in tumors with WGD, we used the timing of the WGD as a reference for timing HLA LOH (**Methods**). We found that breast adenocarcinoma, head and neck squamous cell carcinoma, clear cell renal cell carcinoma, ovarian adenocarcinoma, and leiomyosarcoma tend to have early HLA LOH events (**Fig. S2B**).

### Identification of genetic pathways with immunomodulatory effects

In addition to mutations in the APM, various other pathways have been reported to contribute to immune escape. Existing genomic studies assessing the prevalence of immune escape mechanisms in primary patient tumors suffer from arbitrary selection of a handful of pathways (16). To overcome this limitation, we utilized published genome-wide CRISPR screens that systematically identify the genes involved in different immunomodulatory aspects, such as regulation of MHC-I expression or response to immune-cell-mediated killing (**Fig. 3A**). We identified the top regulators in each study using MAGeCK (26), and conducted Gene Set Enrichment Analysis (GSEA) to identify pathways with immunomodulatory effects. Next, we selected the most frequently enriched immunomodulatory pathways across studies and inferred their mutation timing.

**Figure 3.**
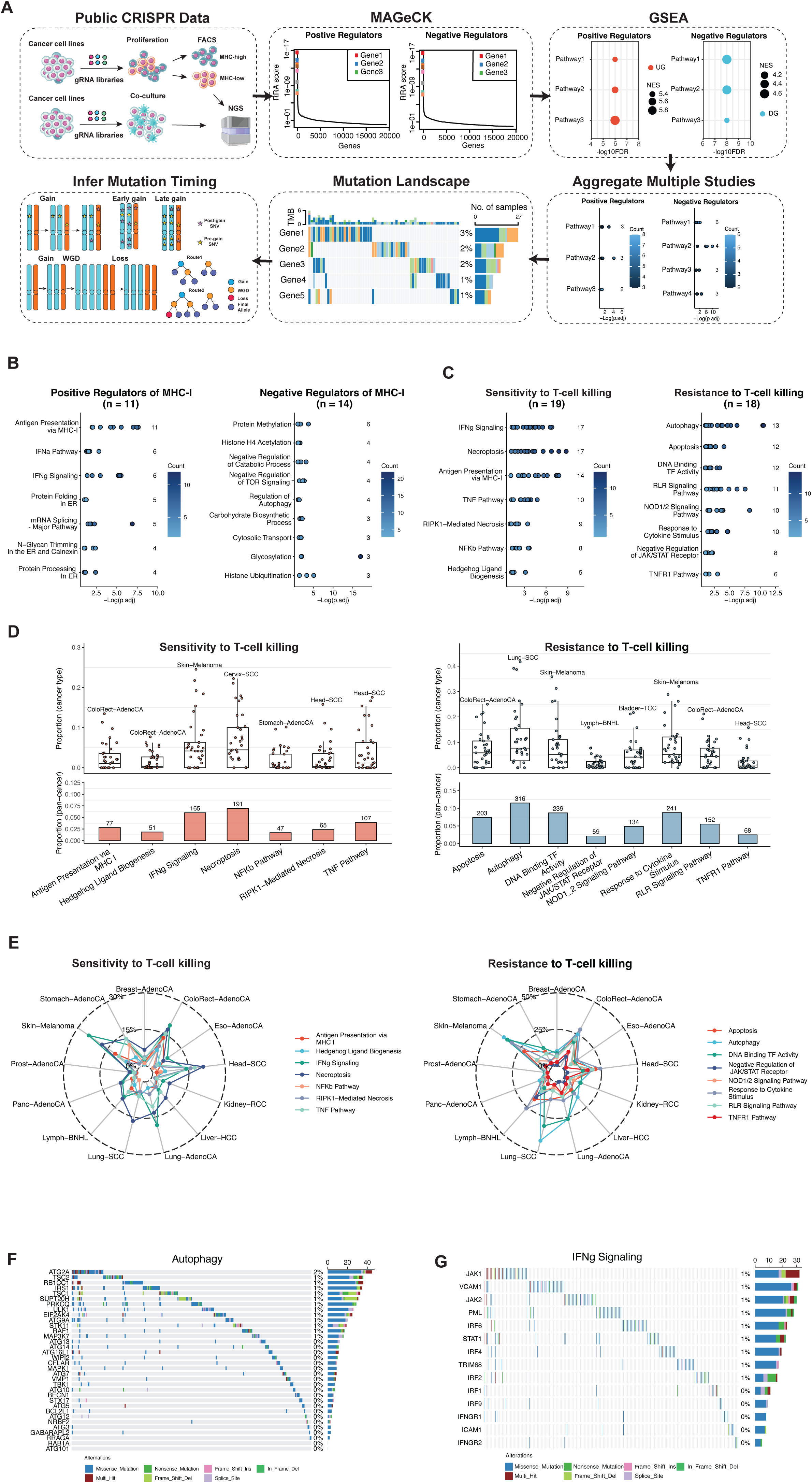
Identification of genetic pathways with immunomodulatory effects. (A) Workflow of identifying CRISPR screen studies for genetic pathways with immunomodulatory effects. (B-C) Summary of CRISPR screen studies reporting the enrichments of MHC-I regulators (B) and T cell killing to tumor cells (C). (D) Prevalence of mutations in the key pathways for T cell killing to tumor cells in all PCAWG cancer samples. (E) Prevalence of mutations in the key pathways for T cell killing to tumor cells in different cancer types. (F-G) Prevalence of point mutations in the representative pathways (autophagy and IFNγ signaling) for the regulation of sensitivity of tumor cells to T cell killing in all PCAWG samples.

Reassuringly, pathways enriched by positive regulators of MHC-I expression included antigen processing and presentation via MHC class I, the IFNα pathway, and IFNγ signaling. Conversely, pathways enriched by negative regulators of MHC-I expression included protein methylation and autophagy, among others (**Fig. 3B**). Regarding the sensitivity of cancer cells to T-cell cytotoxicity, the leading positive regulatory processes were necroptosis, IFNγ signaling, and antigen processing and presentation via MHC class I. The leading negative regulatory processes were autophagy and apoptosis (**Fig. 3C**). Pathways enriched by regulators of response of cancer cells to other immune cells include the PD-L1/PD-1 checkpoint pathway in NK-cell-mediated killing, SWI/SNF in macrophage-mediated killing, and regulation of precursor metabolite generation in γδ T-cell-mediated-killing (**Fig. S3A-C**).

### Mutation frequencies of immunomodulatory pathways across cancer types

Based on the results above, we conducted a pan-cancer analysis of the prevalence of mutations in immunomodulatory pathways. For MHC-I regulators, protein methylation-related genes (*SETD2, KDM6A, MEN1, SETD1B, EZH2, SETDB1*, and *BCOR*) were the most frequently mutated, accounting for 14.6% of all PCAWG samples. Notably, PRC complexes encoded by these genes were identified as negative regulators of MHC-I expression (27) (**Fig. S4A**).

Regarding the sensitivity of cancer cells to T-cell killing, autophagy, as a negative regulator (28), had the highest prevalence of mutations, occurring in 11.5% of PCAWG samples (**Fig. 3D-F**). Mutations in the other representative pathways, IFNγ signaling (29) accounted for 6.0% of all the samples (**Fig. 3G)**. The prevalence of mutations in the regulatory pathways of cancer cells’ responses to other immune-cell-mediated killing varied across cancer types (**Fig. S4B-D**).

We next explored whether mutations in these immunomodulatory pathways were under selection, by calculating the dN/dS ratio. For MHC-I regulators, we observe positive selection in multiple pathways, including the IFNγ pathway, negative regulation of catabolic processes, protein methylation, and regulation of autophagy (**Fig. S5A**). In the regulation of sensitivity of cancer cells to T cell-mediated killing, positive selection was found in DNA Binding transcription factor activity, IFNγ signaling, necroptosis, RIPK1-mediated necrosis, and the TNF pathway (**Fig. 4A-B**). We also observed positive selection of pathways regulating the sensitivity of cancer cells to other immune cell-mediated killing in different cancer types (**Fig. S5B-D**).

**Figure 4.**
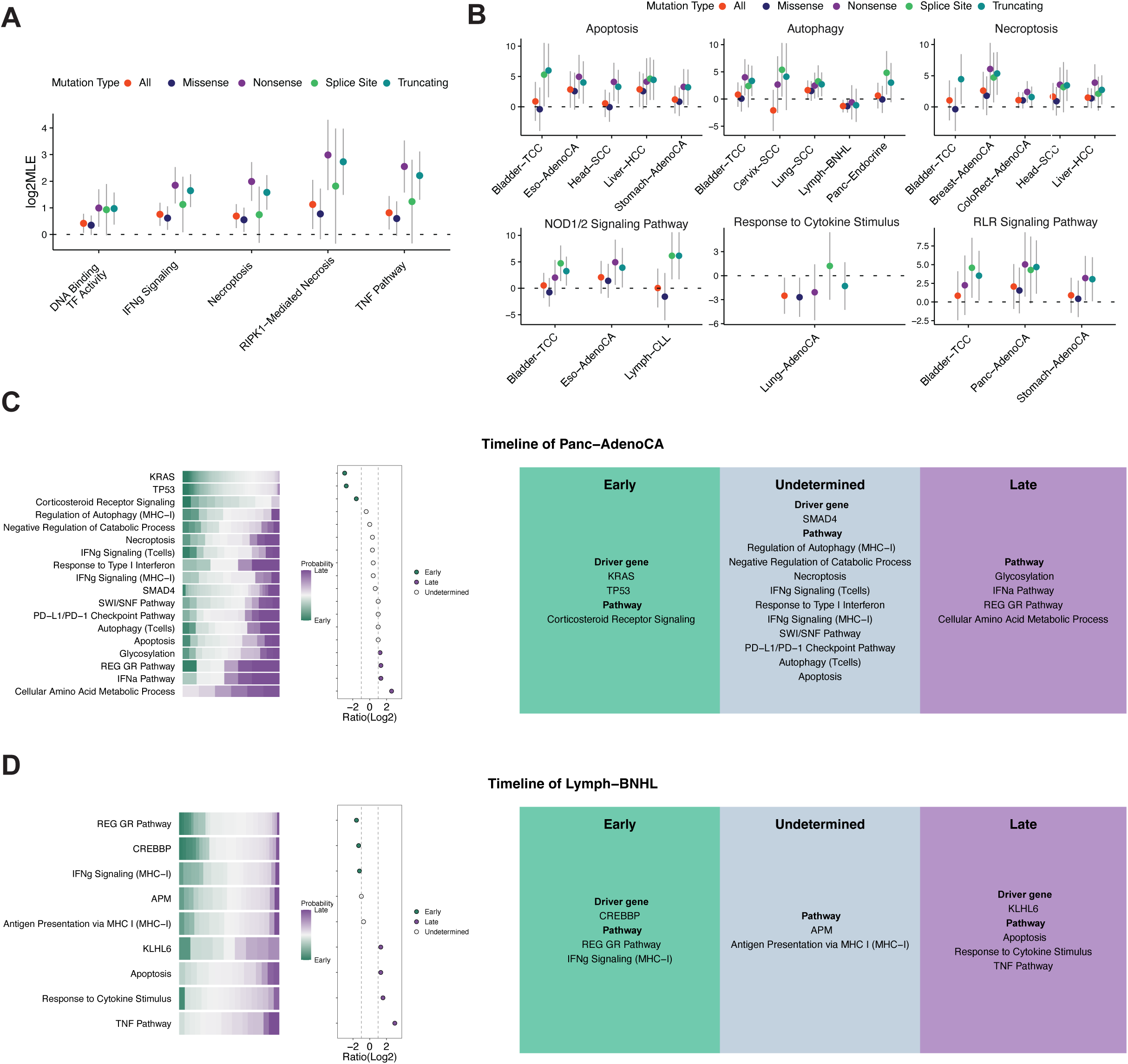
Timing of mutations in genetic pathways with immunomodulatory effects. (A) Positive selection of mutations in regulators for the response of cancer cells to T-cell killing across cancers. (B) Positive selection of mutations in regulators for the response of cancer cells to T-cell killing in each cancer. (C-D) Timing of mutations in genetic pathways with immunomodulatory effects in pancreatic adenocarcinoma (C) and B-cell non-Hodgkin lymphoma (D). Green represents early timing, transitioning gradually to purple for late timing. The color intensity reflects the proportion of early or late events out of 250 samplings, indicating the likelihood of an event occurring early or late. Right Panel: The event is classified as "early" or "late" at the cancer-type level. If the ratio < 0.5, the event is classified as "early”. If the ratio > 2, the event is classified as "late”. Ratio = (Number of late + 1) / (Number of late + 1).

Together, these results reveal a high collective prevalence of mutations in immunomodulatory pathways across cancer types and suggest that different cancers can undergo genomic alterations in different pathways to evade immune surveillance from different immune cells.

### Timing of mutations in immunomodulatory pathways across cancer types

To determine when in cancers’ evolutionary history mutations in immunomodulatory pathways occur, we next applied our GRITIC-based mutation timing approach to identify the timing of mutations in these pathways.

By comparing against the background timing, we found that the timing of mutations in the same pathway varied across cancer types. For regulation of the MHC-I expression, mutations in the protein methylation pathway presented as late events in ovarian adenocarcinoma and bladder transitional cell carcinoma (**Fig. S6A**). For regulation of sensitivity of cancer cells to T cell-mediated killing, mutations in the DNA binding transcription factor activity pathway presented as late events in ovarian adenocarcinoma and breast adenocarcinoma (**Fig. S6B**). We also observed diverse mutation timing of pathways regulating the sensitivity of cancer cells to other cell-mediated killing in different cancer types (**Fig. S6C-D**).

Next, we reconstructed the timeline for mutations in regulators and driver genes with high prevalence in each cancer type. Mutations in the cellular amino acid metabolic process, including *FPGS and GART*, tended to happen late in pancreatic adenocarcinoma (**Fig. 4C**). Mutations in the glucocorticoid receptor regulatory network pathway, including *CREBBP*, tended to occur early, while mutations in the TNF pathways tended to happen late in B-cell non-Hodgkin lymphoma (**Fig. 4D**). Mutations in the NOD1/2 signaling pathway and in DNA binding transcription factor activity, affecting genes such as *MAP3K7, UBE2N, IKBKG, CHUK*, and *IKBKB*, occurred late in breast adenocarcinoma and ovarian adenocarcinoma (**Fig. S7-8**). Mutations in the SWI/SNF pathway were more likely to happen late in hepatocellular carcinoma, with *ARID1A* being one of the most frequently mutated genes in this pathway (**Fig. S9**). The temporal orders of the mutations in the immunomodulatory pathways varied across the other cancer types (**Fig. S10-19**).

Mutations in some immunomodulatory pathways can occur either early or late during cancer development. We wondered whether the timing of such mutations is associated with differences in the tumor microenvironment. Therefore, we compared bulk tumor gene expression between samples with early mutations to those with late mutations for each immunomodulatory pathway in each cancer type. We found that the timing of mutations in immunomodulatory pathways is associated with a different tumor microenvironment in multiple cancer types. For example, patients with early mutations involved in regulation of autophagy had higher expression of negative regulation in immune response in squamous cell lung carcinoma. Patients with late mutations in this pathway had higher expression of genes in leukocyte activity in lung adenocarcinoma. Additionally, patients with late mutations in the apoptosis or DNA binding transcription factor pathway had higher expression of genes involved in leukocyte activation and proliferation in melanoma. Conversely, melanoma patients with early mutations in the autophagy pathway had higher expression of genes involved in leukocyte activity (**Fig. S20**).

Altogether, these results illustrate that different cancer types harness different evolutionary trajectories for immune escape. Revealing these trajectories is critical for early detection and intervention of immune escape and can contribute to immunoprevention of cancers.

## DISCUSSION

“Nothing in biology makes sense except in the light of evolution” - Theodosius Dobzhansky. During the evolution of cancer, cancer cells can develop various strategies to escape immunosurveillance, but the evolutionary process of immune escape is poorly understood, hampering early detection and intervention of immune escape. Answering this question faces two major challenges: (i) identify genes whose mutations can confer immune escape; (ii) determine when such mutations occur in cancer’s evolutionary history. Consequently, previous studies have largely focused on a small number of well-documented pathways in a limited number of cancers.

In this study, we performed data mining of functional genomics screens to systematically identify genes with immunomodulatory effects. We also built upon previous work in evolutionary history inference and built upon our recently developed GRITIC tools to perform fine-scaled timing of events in tumor evolution. Our study reconstructs the evolutionary trajectories of mutations in various immunomodulatory pathways across 38 cancer types in the PCAWG dataset. Together, this study reconstructs the timelines summarizing the typical evolutionary history of immune escape in cancer.

Several limitations in this study should be considered. First, regarding the timing of HLA LOH, as with any statistical approach using multiplicity to infer the evolutionary trajectories, GRITIC cannot quantitatively time chromosomal losses. However, for samples with WGD, we are able to use the relative timing compared to WGD to infer broad timing of HLA LOH. Second, all non-synonymous mutations in a pathway were included for reconstructing the timeline, irrespective of whether they resulted in gain or loss of function. Further investigation into the effects of these mutations on protein function is needed to understand their impact on the evolutionary history of immune escape.

Aligned with previous studies (30,31), we observed that samples with mutations in APM were associated with higher numbers of tumor-infiltrating lymphocytes in several cancer types. Especially in colorectal adenocarcinoma, higher levels of tumor-infiltrating lymphocytes and TMB were found in samples with APM mutations, and APM mutations are under positive selection. These results suggest that the genetic alterations of APM genes might develop escape mechanisms upon strong immune selection pressures.

Besides the diverse timing patterns of APM mutations in cancers, mutations in other immunomodulatory pathways presented different evolutionary trajectories across cancers. The cellular amino acid metabolic process was identified as a negative regulator of tumor cell response to γδT-cell killing, with a mutation prevalence of 2.5% in pancreatic adenocarcinoma. Previous studies reported that γδT cells can infiltrate into the pancreatic adenocarcinoma microenvironment (32,33). Our study suggests that mutations in this pathway can be late-occurring events in pancreatic adenocarcinoma to mediate resistance to γδ-T-cell-mediated killing. The TNF pathway, a positive regulator of tumor cell response to T-cell killing, exhibited mutations in 12.5% of B-cell non-Hodgkin lymphomas. Previous studies revealed that Caspase 8 is often lost or mutated in various tumor types, making it a common target for immune evasion (34,35). Our results suggest that mutations in this pathway happen late in B-cell non-Hodgkin lymphoma.

The DNA-binding transcription factor activity pathway and the NOD1/2 pathway were identified as negative regulators of tumor cell response to T-cell killing, occurring as late events in breast and ovarian adenocarcinomas. The SWI/SNF pathway was identified as a positive regulator of tumor cell sensitivity to macrophage killing, with mutations found in 11.1% of hepatocellular carcinomas. We found that mutations in this pathway are late events in hepatocellular carcinoma evolution. The glucocorticoid receptor regulatory network pathway, a positive regulator of tumor cell response to macrophage killing, showed high mutation prevalence in B-cell non-Hodgkin lymphoma. Our results suggest that mutations in this pathway occur early during B-cell non-Hodgkin lymphoma development, potentially facilitating evasion from macrophage killing. Interestingly, *CREBBP*, a key gene in this pathway, is a known driver gene of B-cell non-Hodgkin lymphoma (36). Consistent with our analysis, it was reported that *CREBBP* can alter tumor-associated macrophage polarization via the FBXW7-NOTCH-CCL2/CSF1 axis, resulting in tumor progression in diffuse large B-cell lymphoma (37). As glucocorticoids are important therapeutic agents for aggressive lymphomas (38), understanding the timing of genetic alterations in this pathway can facilitate more personalized timelines for drug administration.

In conclusion, our study illustrated the evolutionary trajectories of immune escape across cancer types with a broad coverage of immunomodulatory pathways, demonstrating temporally heterogeneous immunogenomic features across cancers. Future studies of time series and multiple-region samples may provide a greater understanding of the evolutionary trajectory of immune escape in cancers.

## METHODS

### Study cohort and data acquisition

Whole genome sequencing, alignment, mutation calling, and copy number data for 2,658 tumors and matched normal pairs across 38 cancer types were obtained from the Pan-Cancer Analysis of Whole Genomes (PCAWG) consortium (39). Thirty-three samples with microsatellite instability status (MSI) were excluded. The VCF file for mutation and copy number variation calling was derived from the ICGC data portal (https://dcc.icgc.org/pcawg).

### Processing the somatic mutation data

The consensus vcf files with somatic mutations were annotated using ANNOVAR. The mutations considered as non-synonymous mutations by the following classifications: “Frame_Shift_Del”, “Frame_Shift_Ins”, “Splice_Site”, “Translation_Start_Site”, “Nonsense_Mutation”, “Nonstop_Mutation”, “In_Frame_Del”, “In_Frame_Ins”, and “Missense_Mutation”. The prevalence of the mutations was visualized by the “Oncoprint” R package.

### Genetic alterations in the antigen presentation machinery

#### HLA somatic mutation calling

In this study, 828 TCGA BAM files were included for the HLA mutation calling and HLA LOH detection based on the GRCh38 version. POLYSOLVER was employed to identify mutations in HLA-A, HLA-B, and HLA-C using its default configuration, without specifying prior population probabilities (40). Initially, HLA allele types for the HLA class I genes were determined, and a winners.hla.txt file was generated for each sample through the POLYSOLVER HLA typing script (shell_call_hla_type). Subsequently, tumor-specific alterations in HLA-aligned sequencing reads were identified by running POLYSOLVER’s mutation-detection script (shell_call_hla_mutations_from_type) on matched tumor–normal pairs, using MuTect. To enhance sensitivity, Strelka (v.2.9.9) was also used to detect short insertions and deletions in HLA-aligned reads, complementing POLYSOLVER’s default caller. Finally, the quality-controlled SNVs and indels were annotated using POLYSOLVER’s annotation script (shell_annotate_hla_mutations). Liftover was performed on the coordinates of HLA-I somatic mutations from the GRCh38 to hg19 genome to make them consistent with the PCAWG consensus data.

#### HLA LOH detection

LOH in *HLA-A, HLA-B, and HLA-C* genes was detected by LOHHLA (10). The same winners.hla.txt files were used as input, with POLYSOLVER’s comprehensive deduplicated FASTA of HLA haplotype sequences as reference. The minimum coverage was set to 5 and maximum mismatch reads were set at 2. The PCAWG consensus purity and ploidy were utilized for the input for LOHHLA. A type-I allele of a patient was annotated as allelic imbalance (AI) if the p-value corresponding to the difference in evidence for the two alleles was <0.01. When the p-value was less than 0.01 and the copy number of the minor allele was less than 0.5, the sample was considered as HLA LOH (10). The coordinates of HLA-I genes were then converted to the hg19 genome to make them consistent with the PCAWG consensus data.

#### Non-HLA somatic mutations of the antigen presentation machinery

The APM compromise the MHC-I complex (*B2M*, *HLA-A*, *HLA-B*, and *HLA-C*), transcription activator (*NLRC5*), immunoproteasome (*PSMB8*, *PSMB9*, and *PSMB10*), peptide clipping (*ERAP1* and *ERAP2*), and antigen loading (*TAP1*, *TAP2*, and *TAPBP*) (19). We also evaluated somatic mutations of the following genes in the APM: *B2M*, *NLRC5*, *PSMB8*, *PSMB9*, *PSMB10*, *ERAP1*, *ERAP2*, *TAP1*, *TAP2*, and *TAPBP*. A sample was defined as an APM mutant if it showed at least one of the following: (1) HLA LOH; or (2) an APM non-synonymous point mutation. Somatic mutations were annotated using ANNOVAR.

### Transcriptional analysis

#### Differentially expressed genes

For the PCAWG-TCGA cohort, we downloaded a raw count matrix for each sample included in PCAWG by “TCGAbiolinks” R package. The gene ID transformations were conducted by the “TransGeneID” function in “MAGeCKflute” R package. The “DESeq2” R package was used to identify differentially expressed genes. Transcripts with a low mean read count (<10) were excluded from the analysis. A binomial Wald test was used after correcting for size factor and dispersion by applying default DESeq2 parameters. To explore significant differences between different groups, we set the threshold of absolute log2FC > 0.5 and adjusted P-value < 0.1.

#### Gene Set Enrichment Analysis

GSEA was performed by the “EnrichAnalyzer” function in the “MAGeCKFlute” R package (41). Gene sets with gene numbers ranging from 5 to 500 were considered. Pathway (KEGG, REACTOME, C2_CP) and C5_BP gene sets were utilized for the analysis. An adjusted p-value of less than 0.1 was considered as the significant level.

#### Immune infiltration

The PCAWG-TCGA RNA-seq data were further utilized to investigate immune cell abundance in different cancer types. Each immune cell type’s relative proportion was estimated for 100 permutations by CIBERSORT with “absolute” mode. LM22 signature matrix was set as the reference in CIBERSORT (42). Quantile normalization of the input mixture was not performed. The absolute mode was applied in CIBERSORT.

### Inference of cancer evolution trajectories

#### Inference of SNV timing

We use GRITIC (18) to quantitatively time clonal copy number gains. GRITIC produces posterior gain timing distributions for clonal copy number segments based on the copy number, tumor purity and the read counts for SNVs in the region of the gain. The segment gain timing distributions were obtained from the original GRITIC study (18).

These gain timing distributions are then used to produce posterior timing distributions for each SNV through a hierarchical sampling process. We first sample a timing state for the corresponding copy number segment and use this to sample a multiplicity state for a given SNV.

This multiplicity state is used to sample a relative time of occurrence between the SNV and the copy number gains and its corresponding segment. The timing of the SNV is then obtained by sampling from a uniform distribution with limits obtained from the gain timing of the preceding and subsequent gains for the SNV. For a posterior sample where no gains occur before the SNV, a lower limit of 0 is used and an upper limit of 1 for a posterior sample where no gains occur after the SNV. We sampled 250 timing values for each SNV. For a posterior sample where an SNV had a subclonal multiplicity state an arbitrary timing value of 1.01 was applied.

The SNV timing is on a scale called mutation time which ranges from 0 to 1, with 0 representing conception and 1 representing the end of the tumor’s clonal evolutionary period.

#### Inference of HLA LOH timing

For the timing of HLA LOH, we assumed that the allele was less likely to lose copy number twice during tumor development with WGD. Therefore, we inferred that if the HLA LOH had a minor copy number equal to 0 and a major copy number greater than 1, the LOH event occurred before the WGD. Subsequently, we assigned a uniform value between 0 and the WGD timing for the HLA LOH.

#### Determining if SNVs occur early or late

We calculated the background timing for each sample by summarizing the mean value for each sampling in the driver genes described in the PCAWG marker paper (39). The average timing of each event in cancer, including mutations in individual driver genes and immunomodulatory pathways, was calculated for each sample. The relative timing for each event was determined by subtracting the background timing from the event’s mean timing. Based on the proportion of the difference, the event in each patient was classified as “early”, “late”, or “undetermined”. If more than 60% of samplings within a patient have a mean timing difference of less than 0, the event is classified as “early”. Conversely, if more than 60% of samplings have a mean timing difference greater than 0, the event is classified as “late”. Events that do not meet either criterion are considered “undetermined” (**Fig. 2A**).

#### Reconstructing the timeline of immune escape in cancer

To construct the timeline for driver genes and immunomodulatory pathways in each cancer type, we selected pathways with mutations in at least five samples that could be determined as occurring early or late in each cancer type. Then, we classified the event as early or late at the cancer-type level based on the ratio between samples with early and late mutations. The ratio was calculated as: (Number of late events + 1) / (Number of early events + 1). If the ratio was less than 0.5, the event was classified as early. Conversely, if the ratio exceeded 2, it was classified as late. Otherwise, we called the event undetermined.

### Identification of core pathways by public CRISPR screening data

#### Data Collection

The published CRISPR screening studies were searched in PubMed based on the keywords “CRISPR”, “Immune” and “Cancer”. The studies with sufficient data regarding MHC regulation and the sensitivity of cancer cells to immune-cell-mediated killing were included, such as T/NK/γδT cell and macrophage. To identify immunomodulatory genes, we collected data from CRISPR screens that identify the regulators of MHC-I expression (27,43–48), response to CD8 T-cell-mediated killing (28,49–56), NK-cell-mediated killing (43,44,50,55,57), macrophage-mediated phagocytosis (58) and γδ T-cell-mediated killing (59).

#### Data Processing

If the study provided the raw FASTQ data, MAGeCK (v0.5.9) was run to generate the count matrix, and the regulators were then obtained by the “RRA” method (26). For those screens using mouse models, the resultant mouse genes were mapped to orthologous human genes using the “TransGeneID” function in “MAGeCKflute” R package, and genes without known homologous relationships were excluded from subsequent analysis. When the raw data was not available, the top regulators for each study were obtained from the tables or supplementary materials in the study. We selected the top 100 genes for negative and positive regulation with immunomodulatory effect for the further analysis.

#### Gene set enrichment for the top regulators

The top 100 positive/negative regulators in each study were respectively unitized for Gene Set Enrichment Analysis. GSEA was performed by the “EnrichAnalyzer” function in the “MAGeCKFlute” R package (41). Gene sets with gene number ranging from 5 to 500 were considered. Pathway (KEGG, REACTOME, C2_CP) and C5_BP gene sets were utilized for the analysis. An adjusted p-value of less than 0.1 was considered as the significant level.

#### Determination of core pathways for immune escape

Next, we selected the featured enrichments for further analysis based on the frequencies of studies reporting these enrichments. For each category of regulation, there can be multiple studies with different top candidates due to the different conditions. To integrate results from multiple studies, we first determined the biological pathways enriched in each study, and then aggregated multiple studies in each category to find the most frequently enriched pathways (**Fig. 3A**). If there was only one study reporting the category for the specific immunomodulatory effect, we selected the pathways based on the most significant p-value and the biological mechanisms. We then combined the immunomodulatory genes in each frequently enriched pathway as a gene set for that pathway and identified their mutations across cancer types in PCAWG.

### Positive selection and tumor mutation burdens

The dNdScv R package was used to quantify selection in somatic evolution including missense, nonsense, and essential splice mutations (21). For truncating mutations, nonsense and essential splice mutations were considered. The local mode was used to analyze the positive selection of the specific gene list. Tumor mutation burdens for each sample were derived from cBioportal (https://www.cbioportal.org/).

### Statistical analysis

All statistical tests were performed in R (4.2.0). Two-sided Wilcoxon tests were used to compare distributions. Tests involving the comparison of proportions in groups were performed using two-sided Fisher’s exact tests. For all statistical tests, all statistical tests were two-sided unless otherwise specified.

## Data Availability

All data utilized in this analysis by PCAWG have been archived in the ICGC data portal (https://dcc.icgc.org/pcawg). Key datasets, including somatic variant calls, subclonal reconstructions, gene expression, and other primary outputs produced by the ICGC/TCGA PCAWG Consortium, are detailed in the PCAWG marker paper (39), and can be downloaded from https://dcc.icgc.org/releases/PCAWG. Further guidance on accessing data, such as raw read files, is available at https://docs.icgc.org/pcawg/data/. Following the data-access policies of the ICGC and TCGA projects, most molecular, clinical, and specimen data are openly accessible without approval. However, access to sensitive information, including germline variants and raw sequencing data, requires researchers to submit applications to the TCGA Data Access Committee (DAC) through dbGaP (https://dbgap.ncbi.nlm.nih.gov/aa/wga.cgi?page=login) for the TCGA portion and to the ICGC Data Access Compliance Office (DACO; http://icgc.org/daco) for the ICGC portion. Additionally, authorization from dbGaP is necessary for researchers to obtain somatic SNVs from TCGA donors.

## Author contributions

S.S.G., P.V.L., and W.C. conceived, designed, and initiated the study. W.C. carried out the majority of the analyses. T.B. and H.A.O. assisted in using GRITIC to infer the timing of genetic events. Z.Z. assisted in processing whole genome sequencing data. W.C., S.S.G., and P.V.L. wrote the manuscript with contributions from all the other authors. S.S.G. and P.V.L. secured funding and supervised the work.

## Acknowledgments

S.S.G. was supported by an NIH Grant (K22CA279077) and the PhRMA Foundation Faculty Starter Grant. P.V.L. is a CPRIT Scholar in Cancer Research and acknowledges CPRIT grant support (RR210006). T.B. was supported by a PhD fellowship from Boehringer Ingelheim Fonds. P.V.L. and T.B. were supported by the Francis Crick Institute which receives its core funding from Cancer Research UK (CC2008), the UK Medical Research Council (CC2008), and the Wellcome Trust (CC2008). The results shown here are in part based upon data generated by the TCGA Research Network: https://www.cancer.gov/tcga.

## Supplementary Figure Legends

**Figure S1. Overview of antigen presentation machinery mutations in PCAWG.** (A) Total number of WGS samples included in the PCAWG dataset. (B) HLA mutation calling by Polysolver and HLA LOH calling by LOHHLA in PCAWG-TCGA samples. (C) dN/dS ratios for antigen presentation machinery mutations in each PCAWG cancer type.

**Figure S2. Temporal order of antigen presentation machinery mutations and key events in cancers.** (A) Distribution of timing of antigen presentation machinery point mutation across cancer types. The plots show the mean timing of 250 timing samples for all APM mutations in each sample. (B) Distribution of timing of HLA LOH, and WGD across cancer types.

**Figure S3. Determining genetic pathways with immunomodulatory effects.** (A) Summary of CRISPR screen studies reporting the enrichments of regulators for the response of cancer cells to NK-cell killing. (B) Summary of CRISPR screen studies reporting the enrichments of regulators for the response of cancer cells to macrophage killing. (C) Summary of CRISPR screen studies reporting the enrichments of regulators for the response of cancer cells to γδ T-cell killing.

**Figure S4. Prevalence of mutations in genetic pathways with immunomodulatory effects.**

(A) Prevalence of mutations in the key pathway for regulators of MHC-I regulation. (B) Prevalence of mutations in the key pathway for regulators for the response of cancer cells to NK-cell-mediated killing. (C) Prevalence of mutations in the key pathway for regulators for the response of cancer cells to macrophage-mediated killing. (D) Prevalence of mutations in the key pathway for regulators for the response of cancer cells to γδT-cell-mediated killing. Top panels show the prevalence of mutations in the key pathways for T cell killing to tumor cells in all PCAWG cancer samples. Bottom panels show the prevalence of mutations in the key pathways for T cell killing to tumor cells in different cancer types.

**Figure S5. Positive selection of mutations in genetic pathways with immunomodulatory effects.** (A) Positive selection of mutations in regulators for MHC-I expression. (B) Positive selection of mutations in regulators for the response of cancer cells to NK-cell-mediated killing. (C) Positive selection of mutations in regulators for the response of cancer cells to macrophage-mediated killing. (D) Positive selection of mutations in regulators for the response of cancer cells to γδT-cell-mediated killing.

**Figure S6. Timing of mutations in the pathways for regulating MHC-I expression and the response of cancer cells to T-cell-mediated killing.** (A) Timing of mutations in regulators for MHC-I expression. (B) Timing of mutations in regulators for the response of cancer cells to T-cell-mediated killing. (C) Timing of mutations in regulators for the response of cancer cells to NK-cell-mediated killing. (D) Timing of mutations in regulators for the response of cancer cells to macrophage-mediated killing. Green represents early timing, transitioning gradually to purple for late timing. The color intensity reflects the proportion of early or late events out of 250 samplings for each patient, indicating the likelihood of an event occurring early or late.

**Figure S7-S19. Timeline of mutations in the immunomodulatory pathways and key events in each cancer type.** Green represents early timing, transitioning gradually to purple for late timing. The color intensity reflects the proportion of early or late events out of 250 samplings, indicating the likelihood of an event occurring early or late. Right Panel: The event is classified as early or late at the cancer-type level. If the ratio < 0.5, the event is classified as early. If the ratio > 2, the event is classified as late. Ratio = (Number of late + 1) / (Number of late + 1). Otherwise, we called the event undetermined.

**Figure S20. Enrichments of differentially expressed genes by the SNV timing in genetic pathways with immunomodulatory effects in multiple cancer types.** The “higher-in-late-events” enrichments indicate that these genes were more highly expressed in patients with later events compared to those with earlier events. Conversely, the “higher-in-early-events” enrichment shows that the expression of these genes was greater in patients with early events. Red dots represent enrichments that are upregulated in patients with late events, while the blue dot represents enrichment that is up regulated in patients with early events.

