## Supplementary figures and images for "Evolutionary trajectories of immune escape across cancers"

### SFigure1.pdf

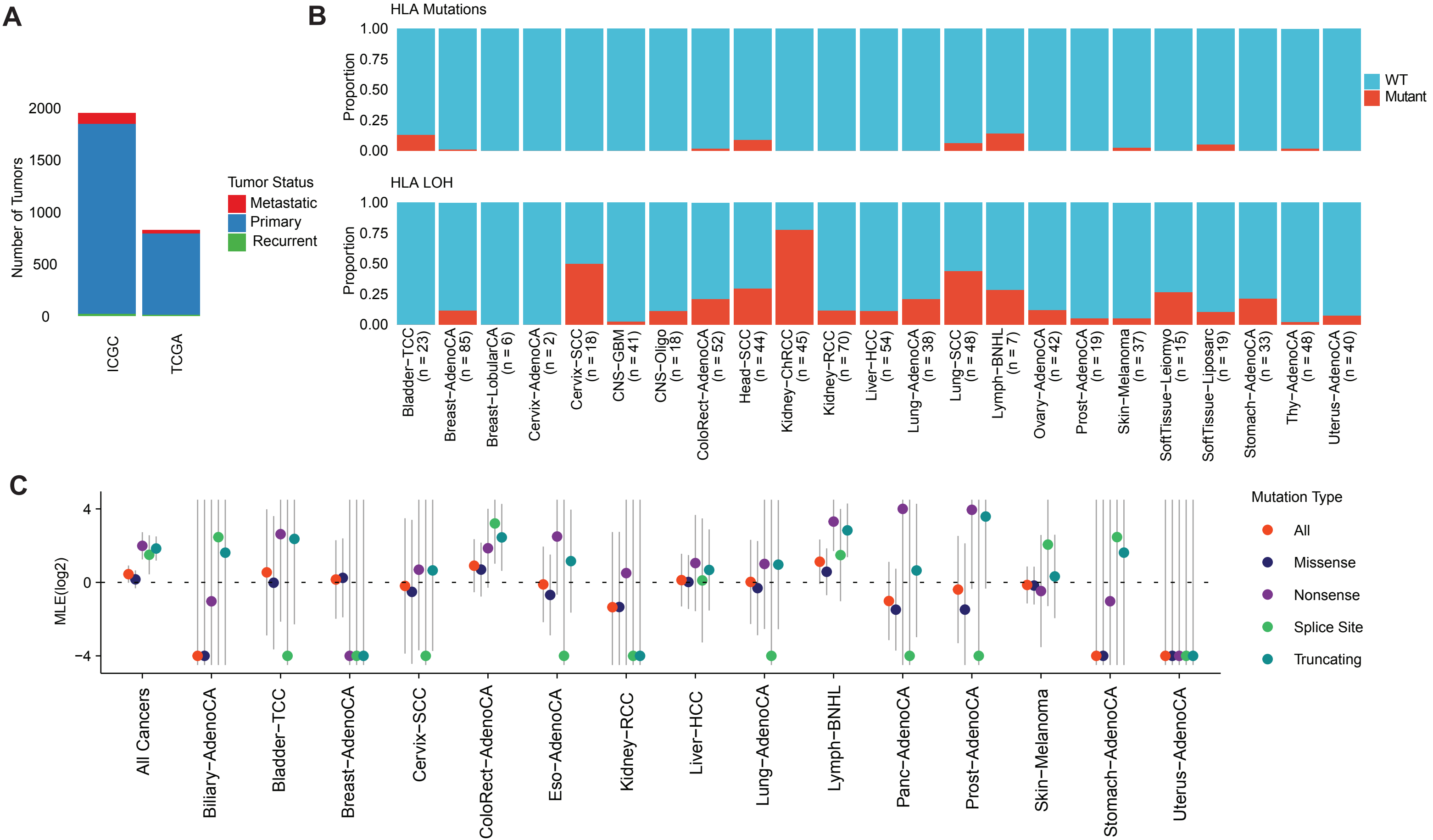

### SFigure2.pdf

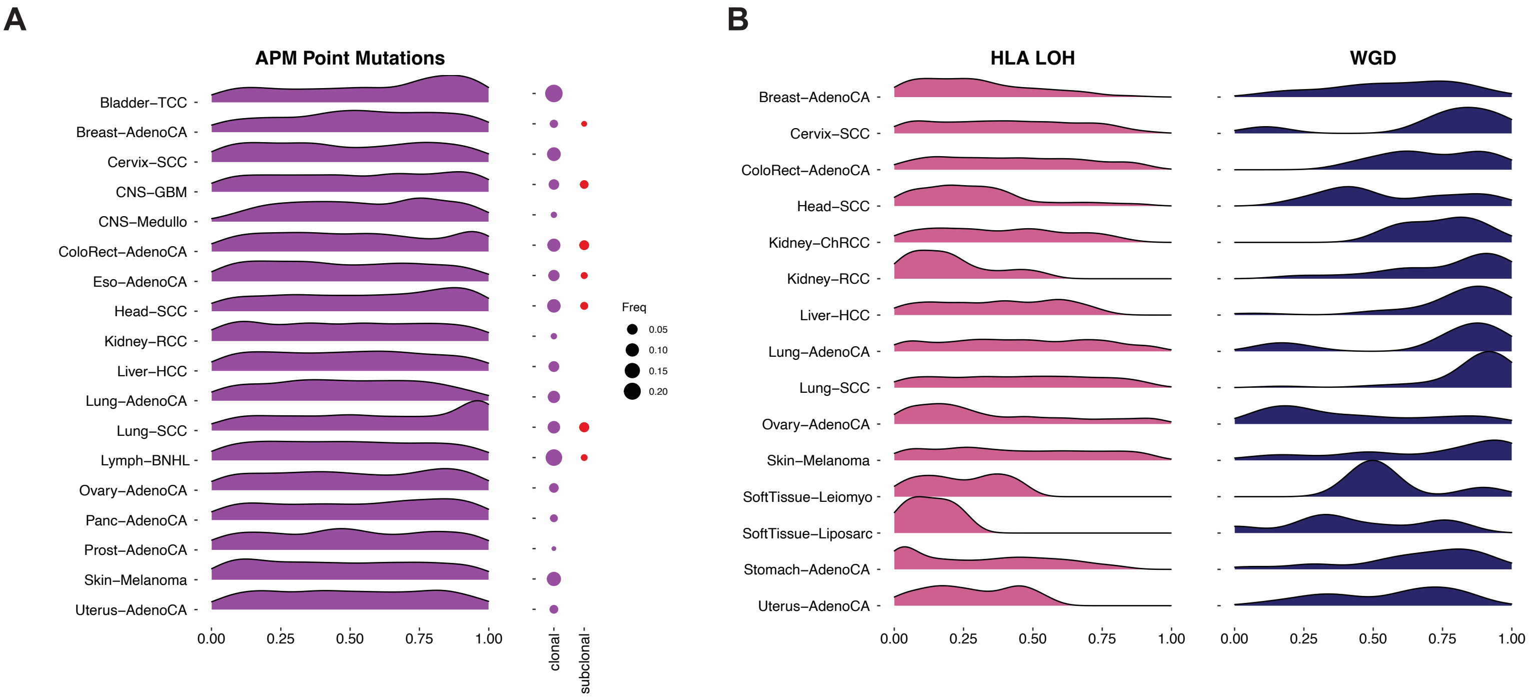

### SFigure3.pdf

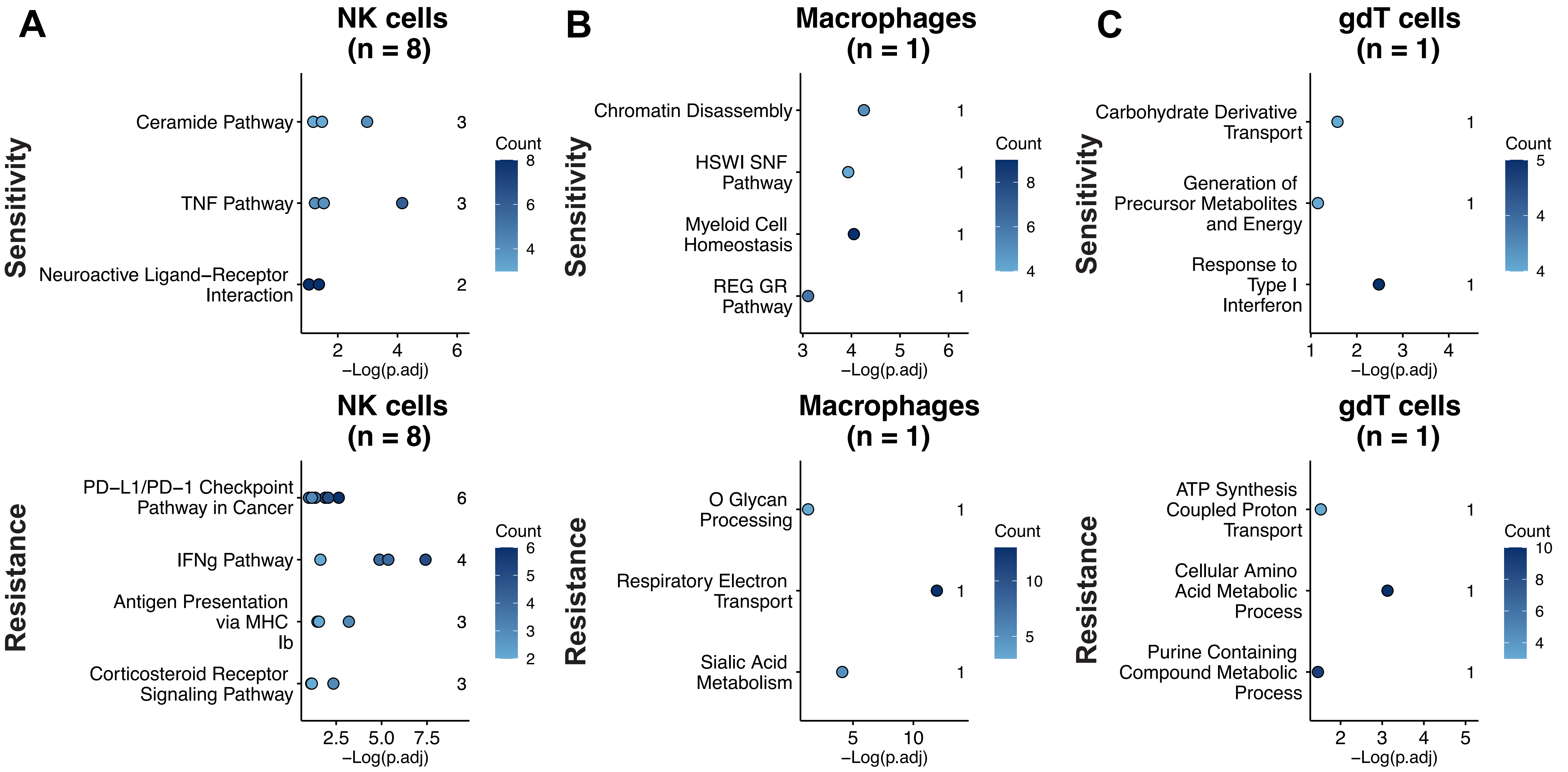

### SFigure4.pdf

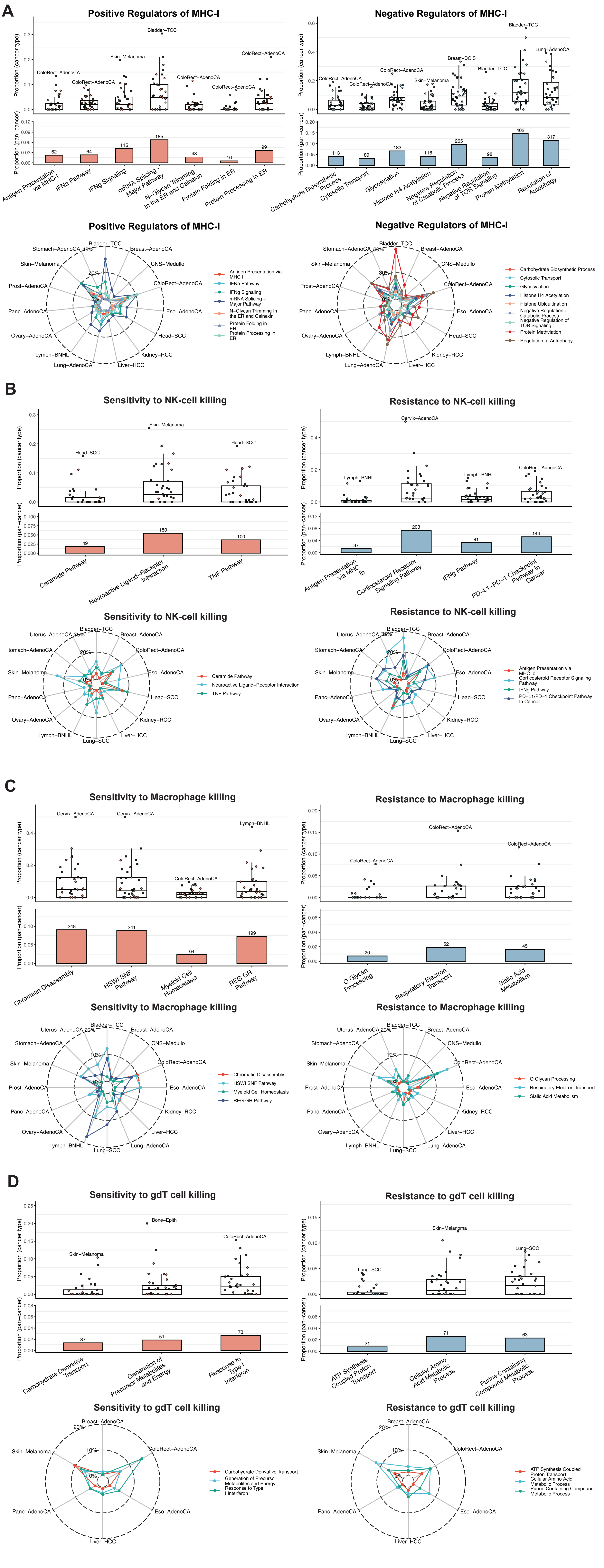

### SFigure5.pdf

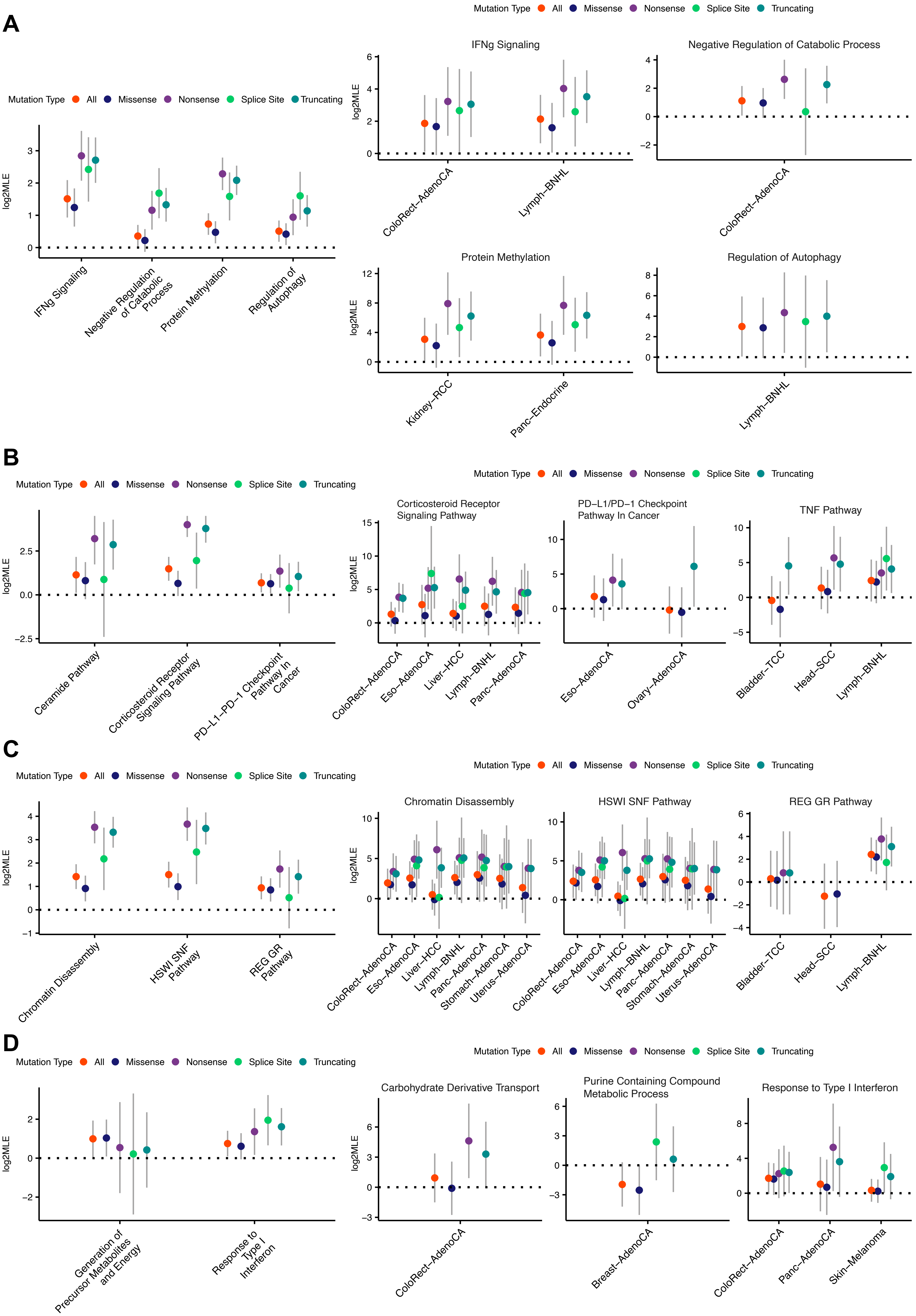

### SFigure6.pdf

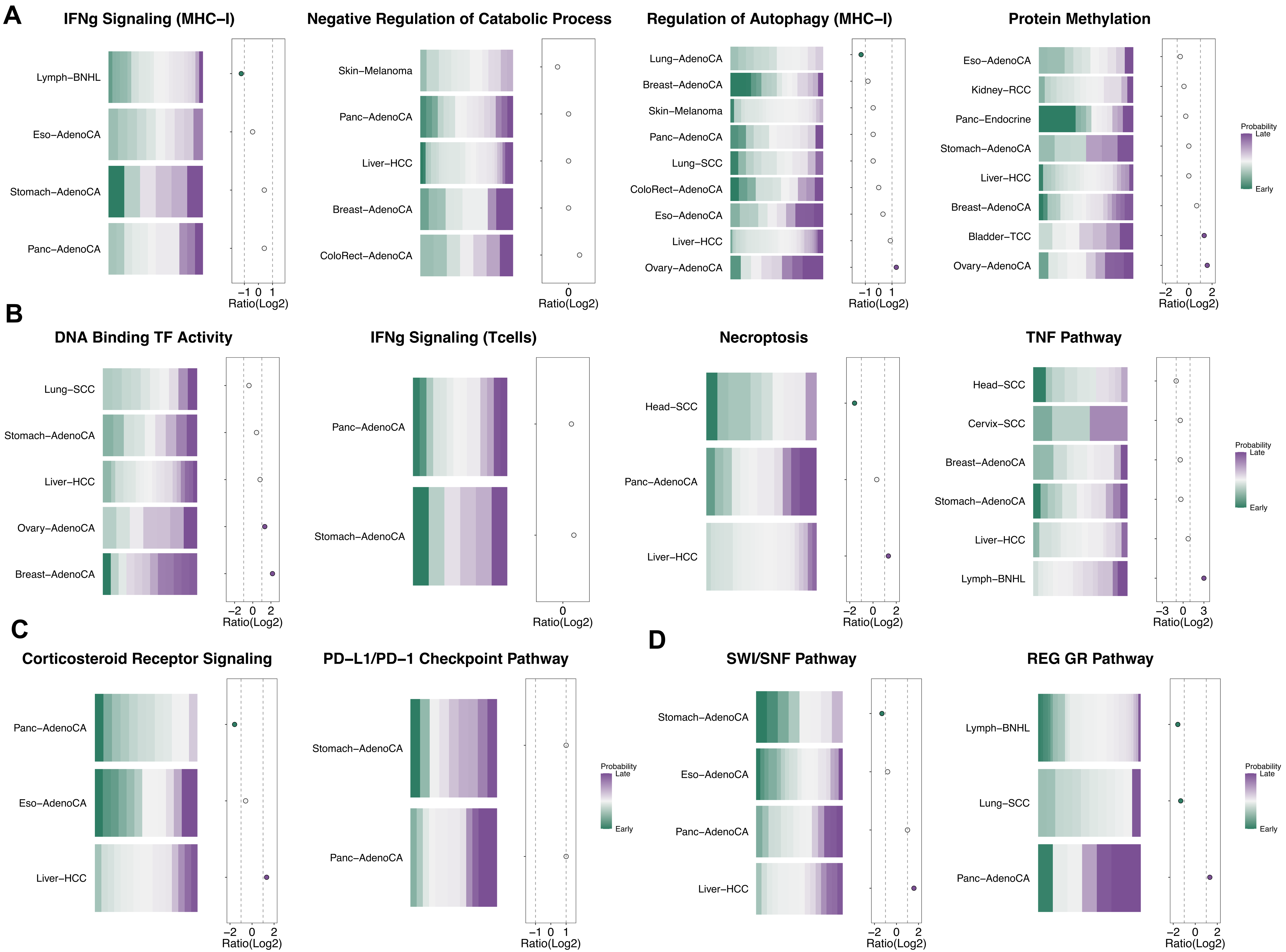

### SFigure7_Breast-AdenoCA.pdf

#### Timeline of Breast–AdenoCA

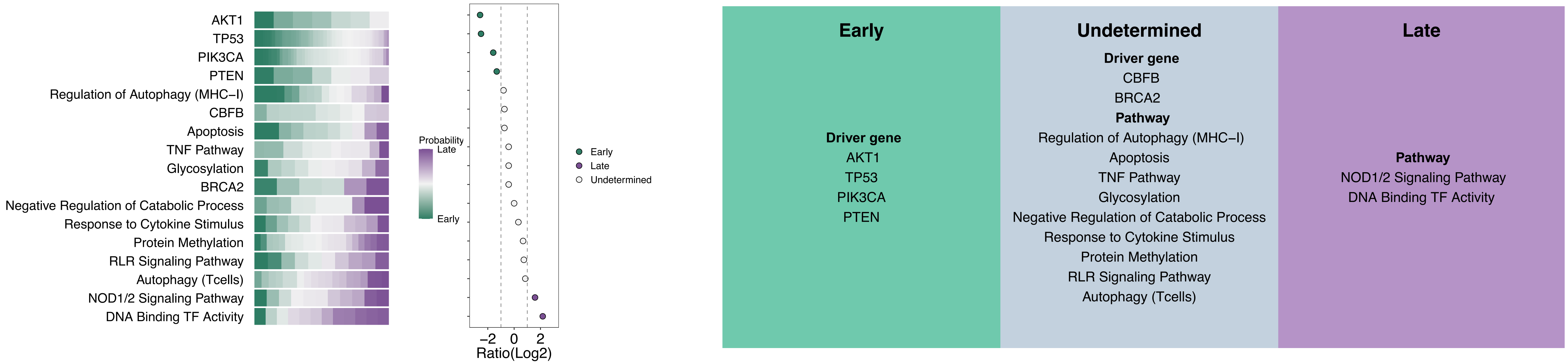

### SFigure8_Ovary-AdenoCA.pdf

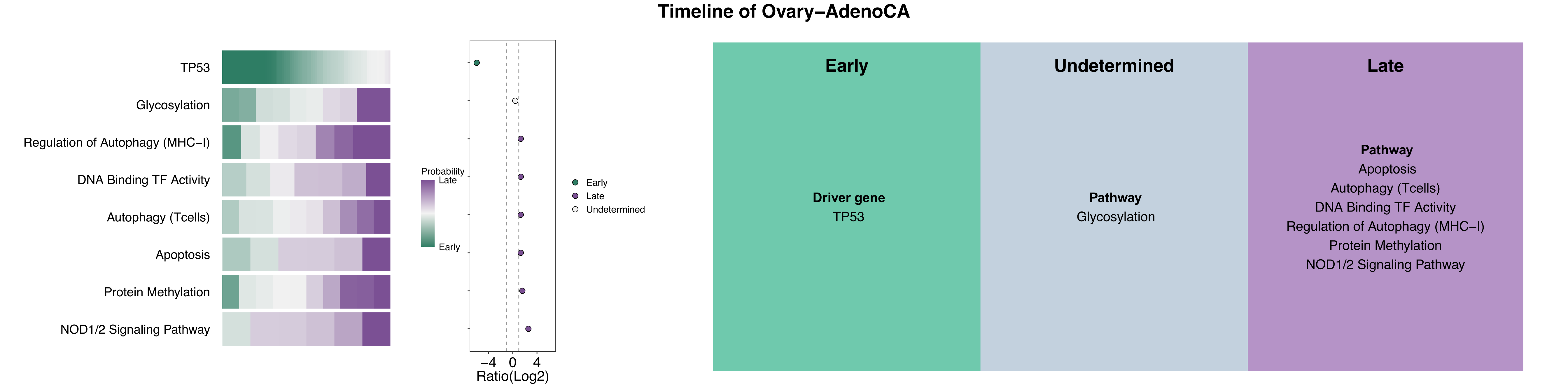

### SFigure9_Liver-HCC.pdf

##### Timeline of Liver–HCC

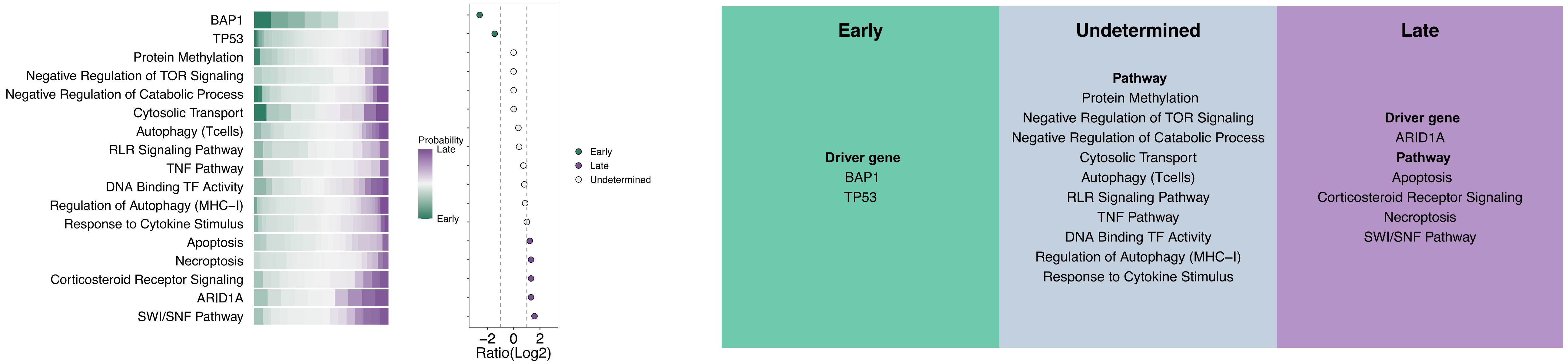

### SFigure10_Bladder-TCC.pdf

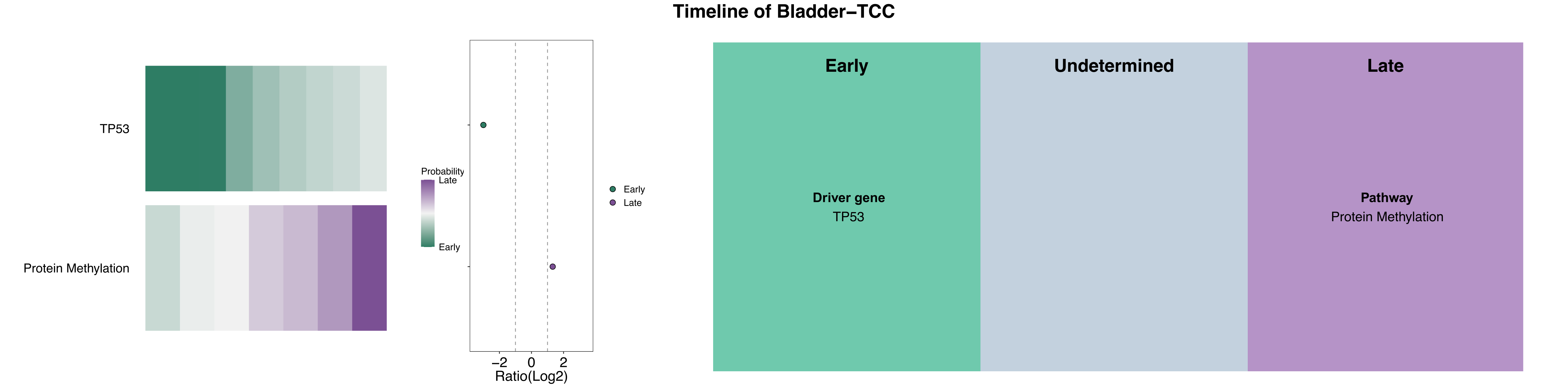

### SFigure11_ColoRect-AdenoCA.pdf

#### Timeline of ColoRect–AdenoCA

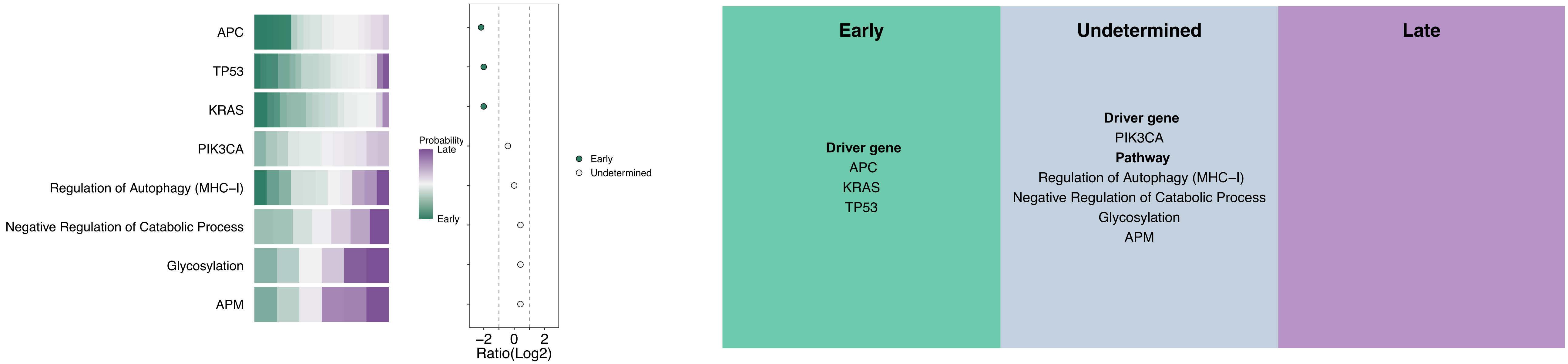

### SFigure12_Eso-AdenoCA.pdf

#### Timeline of Eso−AdenoCA

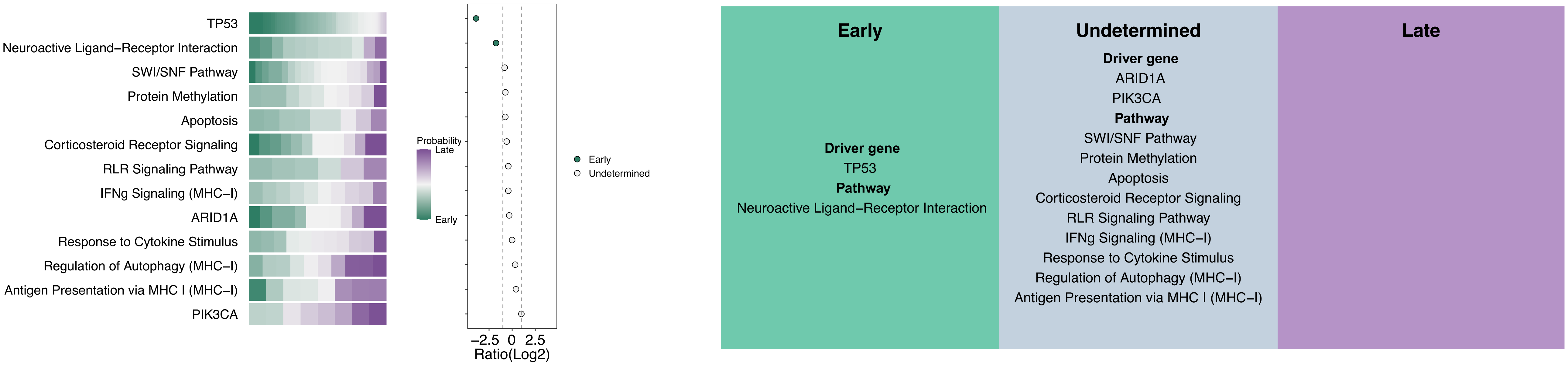

### SFigure13_Head-SCC.pdf

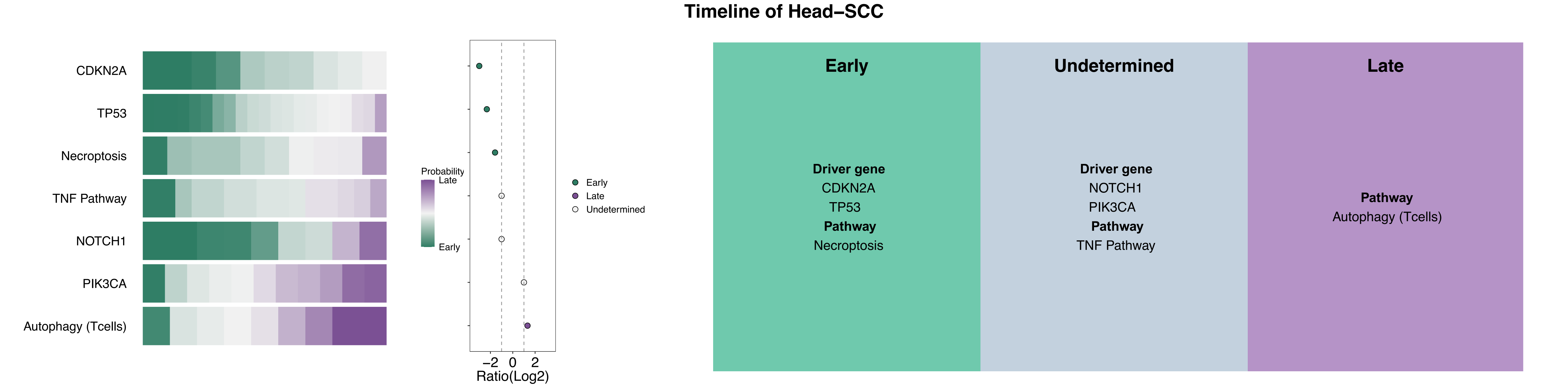

### SFigure14_Kidney-RCC.pdf

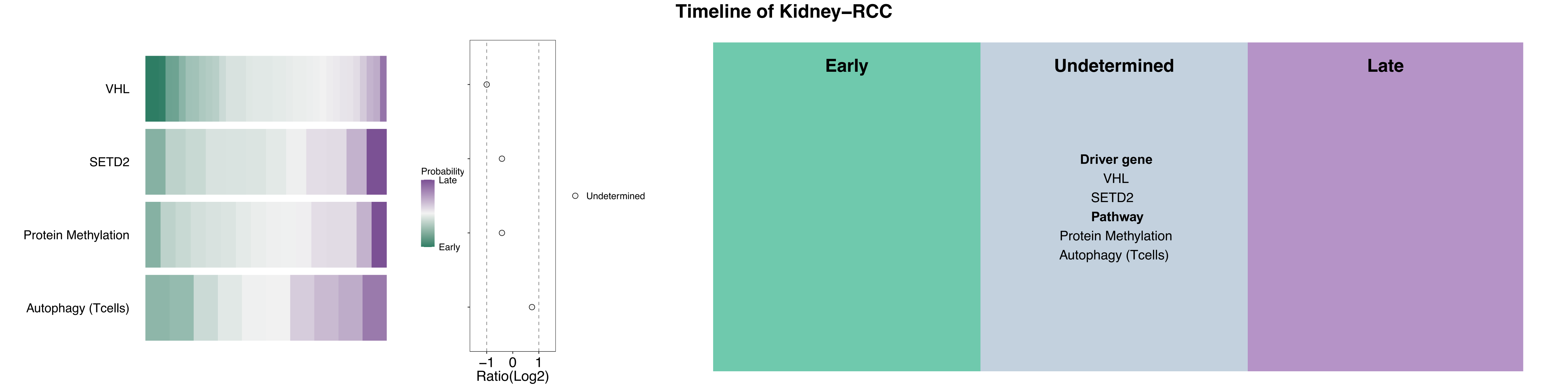

### SFigure15_Lung-SCC.pdf

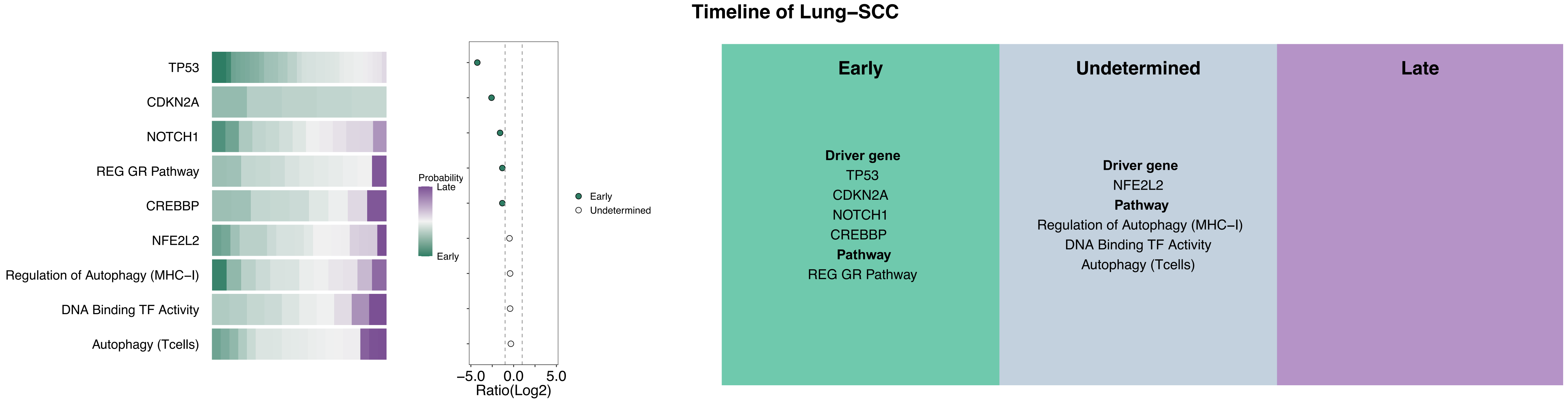

### SFigure16_Panc-Endocrine.pdf

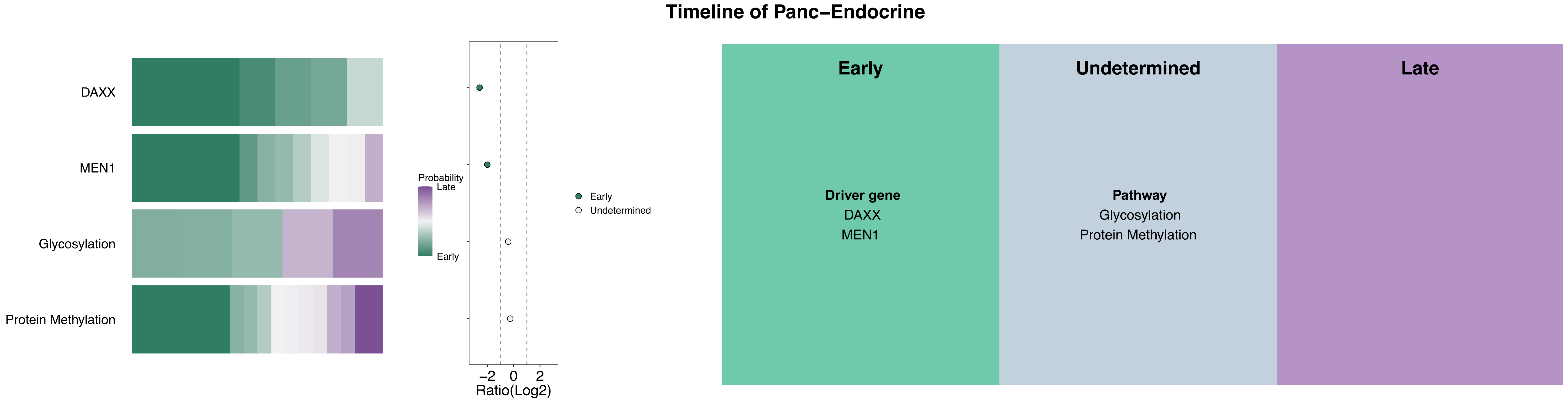

### SFigure17_Skin-Melanoma.pdf

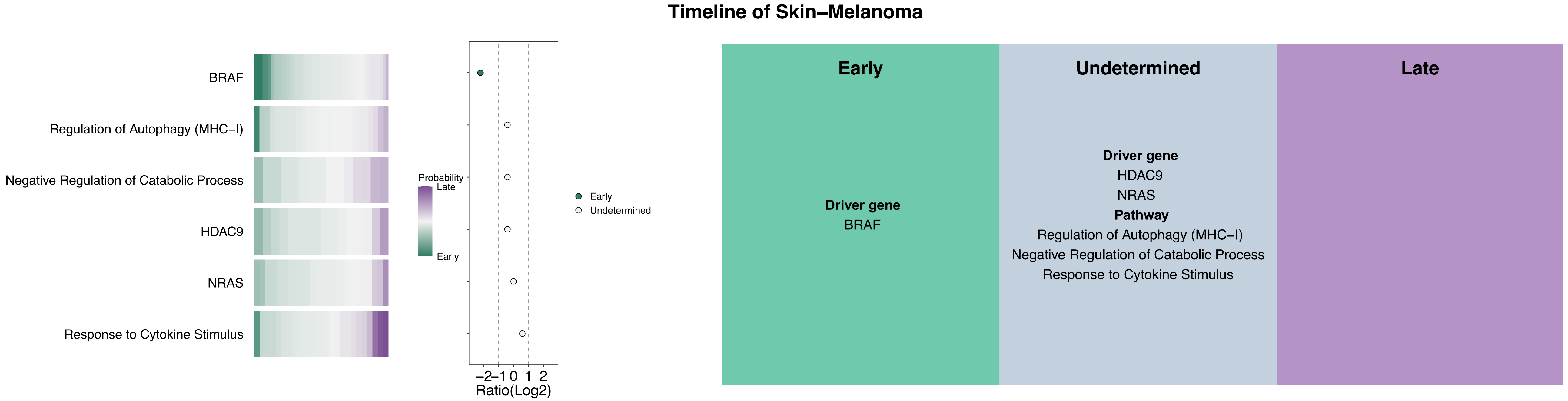

### SFigure18_Stomach-AdenoCA.pdf

#### Timeline of Stomach–AdenoCA

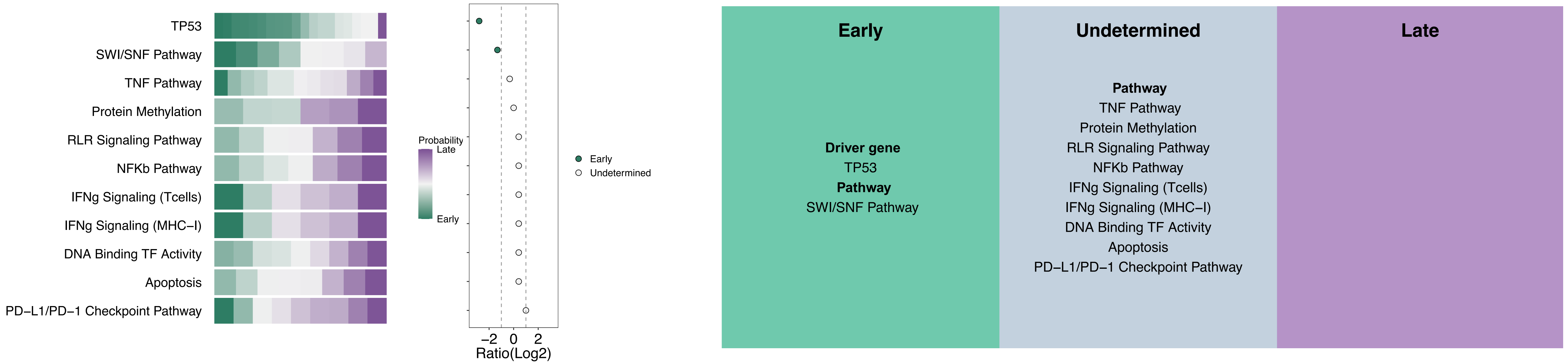

### SFigure19_Uterus-AdenoCA.pdf

#### Timeline of Uterus–AdenoCA

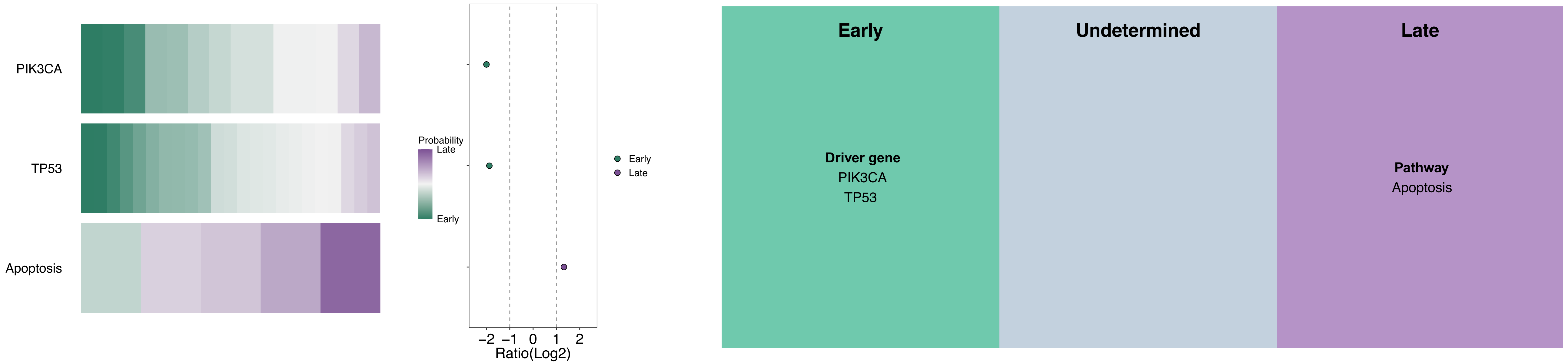

### SFigure20.pdf

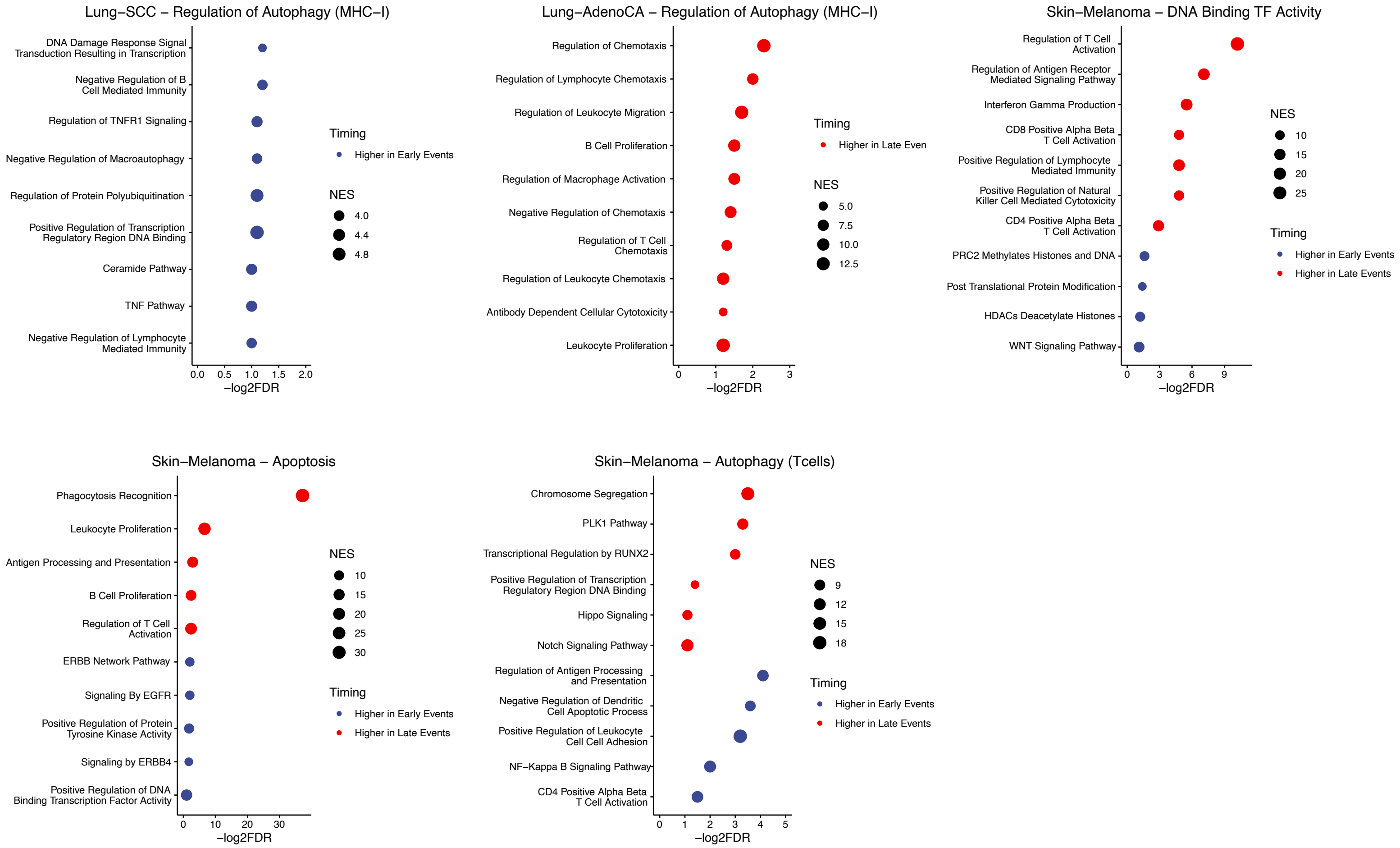
